# ST-Analyzer: A Packaged Web and Command-Line Interface for Simulation Trajectory Analysis

**DOI:** 10.64898/2026.02.06.704471

**Authors:** Nathan R. Kern, Soohyung Park, Yiwei Cao, Wonpil Im

## Abstract

As high-performance computing provides the ability to generate and analyze ever larger simulation trajectories, the challenges in learning, applying, and sharing the best analytical practices become more salient. Extracting reproducible scientific insights from simulation requires a thorough understanding of many computing topics unrelated to the molecular systems being modeled and simulated. While the rapid development of the technologies used for analysis makes previously impossible studies into routine work, the growing repertoire of software combined with the specificity of the ecosystems that they rely on can easily break the programs used in older studies. In this work, we present ST-Analyzer, a simulation trajectory analysis suite with command-line (CLI) and graphical (GUI) user interfaces. ST-Analyzer is distributed freely as an open-source conda-forge package with support for macOS, Linux, and Windows (via WSL2). Besides facilitating several common analysis tasks, the GUI shows users the exact commands necessary to repeat the same tasks on the command-line. We demonstrate ST-Analyzer’s capabilities by reproducing several results from previously published simulation studies on the lipid parameters of heterogeneous biomembranes and the behavior of a SARS-CoV-2 spike protein-antibody complex. We expect ST-Analyzer to be useful to experts for quickly setting up common analysis tasks and to nonexperts as a guided introduction to simulation analysis using both GUI and CLI. ST-Analyzer is freely available at https://github.com/nk53/stanalyzer.

## Introduction

Molecular dynamics (MD) simulation has become a useful supplement to the experimental study of biomolecules at the atomic scale. The level of detail achievable by simulation programs is limited primarily by the resolution and accuracy of the force fields, the validity of the constructed molecular models, the computational power of the high-performance computing (HPC) centers running and analyzing simulations, and the incredible volume of space needed to store simulation data. Currently, medium-scale HPCs available to university researchers can easily generate terabytes of simulation data per week. Although technologies and techniques are improving, both experts and nonexperts struggle to obtain reproducible insights from this glut of data through MD simulation trajectory analysis.

A variety of tools are available to academic researchers to facilitate useful data extraction using well-tested techniques. This includes standalone programs like Qhull^1^; robust, extensible software suites for modelling and analysis like GROMACS,^2^ VMD,^3^ and CHARMM;^4^ and numerous programming libraries with permissive licensing such as MDAnalysis,^5^ MDTraj,^6^ CPPTraj,^7^ LOOS/PyLOOS,^8^ Numpy,^9^ Scipy,^10^ etc. It is common to use many such tools to analyze the same simulation trajectory. A crucial decision in analysis is whether enough high-quality data have been collected to justify stopping a simulation and releasing its associated computational resources. This decision requires periodic evaluations of trajectories. Researchers with command-line knowledge benefit greatly from tools that expose their different capabilities through a command-line interface (CLI), as they facilitate the creation of custom analysis pipelines with (relatively) simple shell scripting languages like bash.

While programming libraries enable the development and publication of custom analyses, they also require researchers to understand programming conventions, data-structure intricacies, and the peculiarities of each library’s application programming interface (API). Likewise, each software suite’s CLI has its own conventions that the researcher must learn to use the tool effectively. Thus, these tools have a steep learning curve; even a standard task like computing a protein backbone’s root-mean-square deviation (RMSD) requires a familiarity with filesystems, command-line syntax, and several file formats. The most powerful tools also expose a daunting number of options that, if incorrectly specified, can result in subtle but significant errors in the final analysis. The result is that researchers who are new to MD simulation spend a disproportionate amount of time troubleshooting software instead of generating and interpreting results.

CHARMM-GUI is a cyberinfrastructure for building simulation-ready molecular models and input files for most MD engines.^11–13^ Although it uses CHARMM extensively under the hood, a core philosophy of CHARMM-GUI is that its outputs should be usable by other popular simulation programs with minimal modification. Although most CHARMM-GUI input generators^14^ have extensive options, it aims to expose those options in a user-friendly way and provide sensible default values where possible. Nonetheless, CHARMM-GUI lacks similar workflows to run and analyze simulations.

In this work, we present ST-Analyzer (https://github.com/nk53/stanalyzer), which aims to bridge the gap between running a simulation and generating initial results. While ST-Analyzer was previously published in 2014,^15^ this work is a complete refactor. The main design differences emphasize ease of installation, cross-platform usability, and minimizing the extent of knowledge of ST-Analyzer internals necessary to extend it with new analyses. To this end, we have contributed ST-Analyzer as a new recipe with permissive licensing for the well-known package manager conda in the conda-forge repository.^16^ The conda-forge framework facilitates software development, testing, and distribution via continuous integration services and global distribution servers.

Although publication-ready results often include bespoke analysis and visualization procedures, the extent of customization necessary tends to become clear only after performing a standard set of analyses, such as measuring the RMSD of specific protein segments, sterol tilt angles, area per lipid, etc. Furthermore, by providing a CLI tool with a locally hosted web GUI that can be installed with conda, we hope to bridge the transition that many researchers in MD simulation encounter when interacting with HPCs: moving from GUIs to using the CLI to perform analysis.

### Software Design and Implementation

ST-Analyzer consists of 4 components: (i) a CLI utility *stanalyzer* for running analysis routines, (ii) self-contained analysis procedures, (iii) a web GUI that facilitates project configuration and invocation of the CLI tool, and (iv) a server program that tracks the status of analyses requested through the GUI and serves pages to the user’s preferred web browser.

### Command-line interface: *stanalyzer*

The *stanalyzer* command-line program locates the desired analysis procedure, reads options from the command-line, and passes them to the analysis’s main function. **Figure 1** shows a usage example, illustrating that even the simplest possible analysis (reading box dimensions from each simulation frame) takes several options. While *stanalyzer* can be run by specifying each of these options manually, we anticipate that some options will rarely change between runs, e.g., the structure and coordinate files to use for analysis. To reduce the number of options that need to be given manually, *stanalyzer* looks for a config file that stores defaults for the most commonly used settings. This file can be created in the web GUI or interactively on the command-line with the *stanalyzer* config helper tool. As a simple JSON file, the settings are human-readable and easily changed in a text editor (**Figure S1** and SI **S1. Schema for** `analysis.yml`

**Figure 1.**
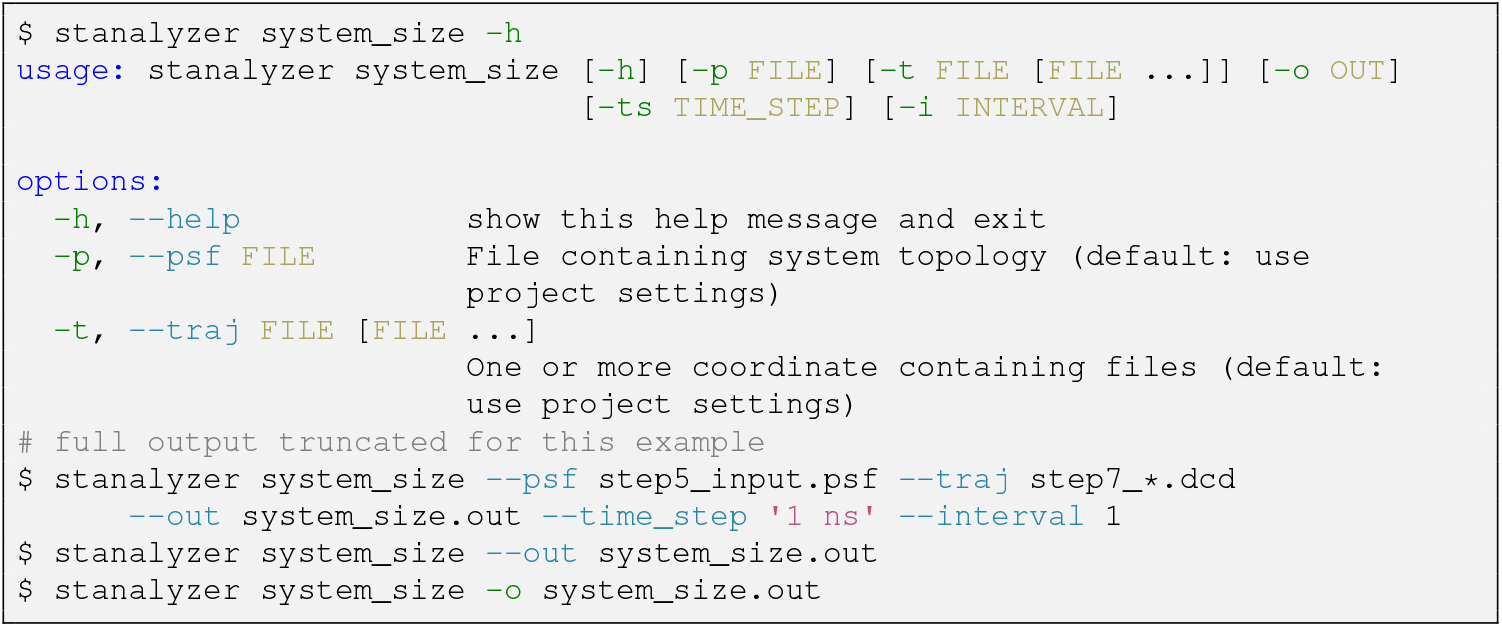
Command-line *stanalyzer* usage example.

### Analysis programs

All analysis programs are located in the Python package’s analysis subdirectory. The minimum requirements to make a new analysis routine available to the CLI are to (i) be placed in ST-Analyzer’s analysis subdirectory, (ii) define a nullary *get_parser()* function that returns an *argparse*.*ArgumentParser* object, and (iii) define a unary main(settings) function that accepts a *dict* of settings. **Figure S2** shows a minimal script to wrap an external program that handles its own arguments. This allows non-Python programs to be used with ST-Analyzer’s CLI. The list of implemented analyses is summarized in **Table 1**. To expose an analysis to the web server, a future developer must describe the web form according to the YAML format shown in **Figure S3** and SI **S2. Schema for** `analysis.yml`.

**Table 1.**
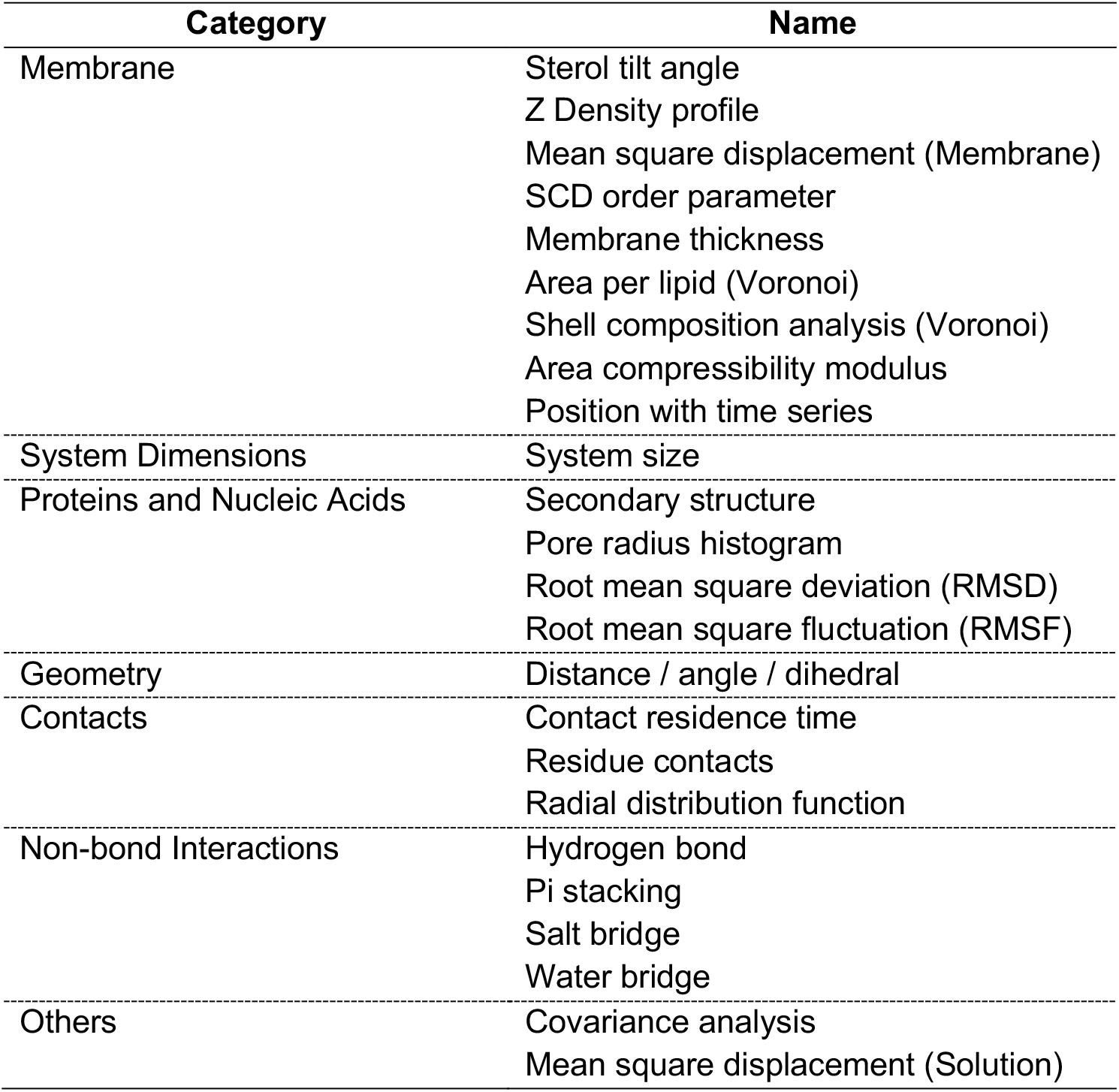
Available analysis programs.

### Web server

The web server uses the FastAPI^17^ framework to serve web pages, validate user input, manage software settings, and track the status of analysis jobs created through the web GUI (see the next subsection). Users can start the self-hosted server by invoking the *sta-server* command with no options. This opens a debug log and displays the address from which users can connect to the server. The program is designed to be runnable from a personal computer and has been tested on macOS, Linux, and the Windows Subsystem for Linux v2 (WSL2).

By default, it only accepts connections from localhost, but the underlying server program (uvicorn) can be configured to accept remote connections over transport layer security (TLS). Remote connection management and authentication can also be delegated to a proxy such as Nginx.^18^ To aid in debugging and to help train researchers to use the command line, the web server records the exact command used to run each analysis in the project’s output directory. The arguments and PID of the corresponding *stanalyzer* task are stored in a SQLite database. When a user requests to check a PID’s status, the server checks if the process still exists and returns a message either saying that the job is still running or containing the contents of its output and (if applicable) error logs.

### Web GUI

The web GUI uses responsive design principles, allowing it to adapt smoothly to diverse screen sizes and changes in window size. After launching the web server with default settings, the GUI is accessed by entering the URL http://127.0.0.1:8000 into a web browser’s address bar. The user must first configure a project, which tells ST-Analyzer where to find the input structure and trajectory files and where to write analysis output (**Figure 2**). Next, the user selects one or more analyses to perform. Each analysis defines its own web form (see **Figure S3**) to select options. On form submission, the web server translates the form into a set of command-line options and responds with the exact commands used to start each requested analysis. If an analysis fails to start, the response instead contains the cause of failure, e.g., an input validation error. Either way, the response is displayed to the user in a non-modal dialog (**Figure 2C**).

**Figure 2.**
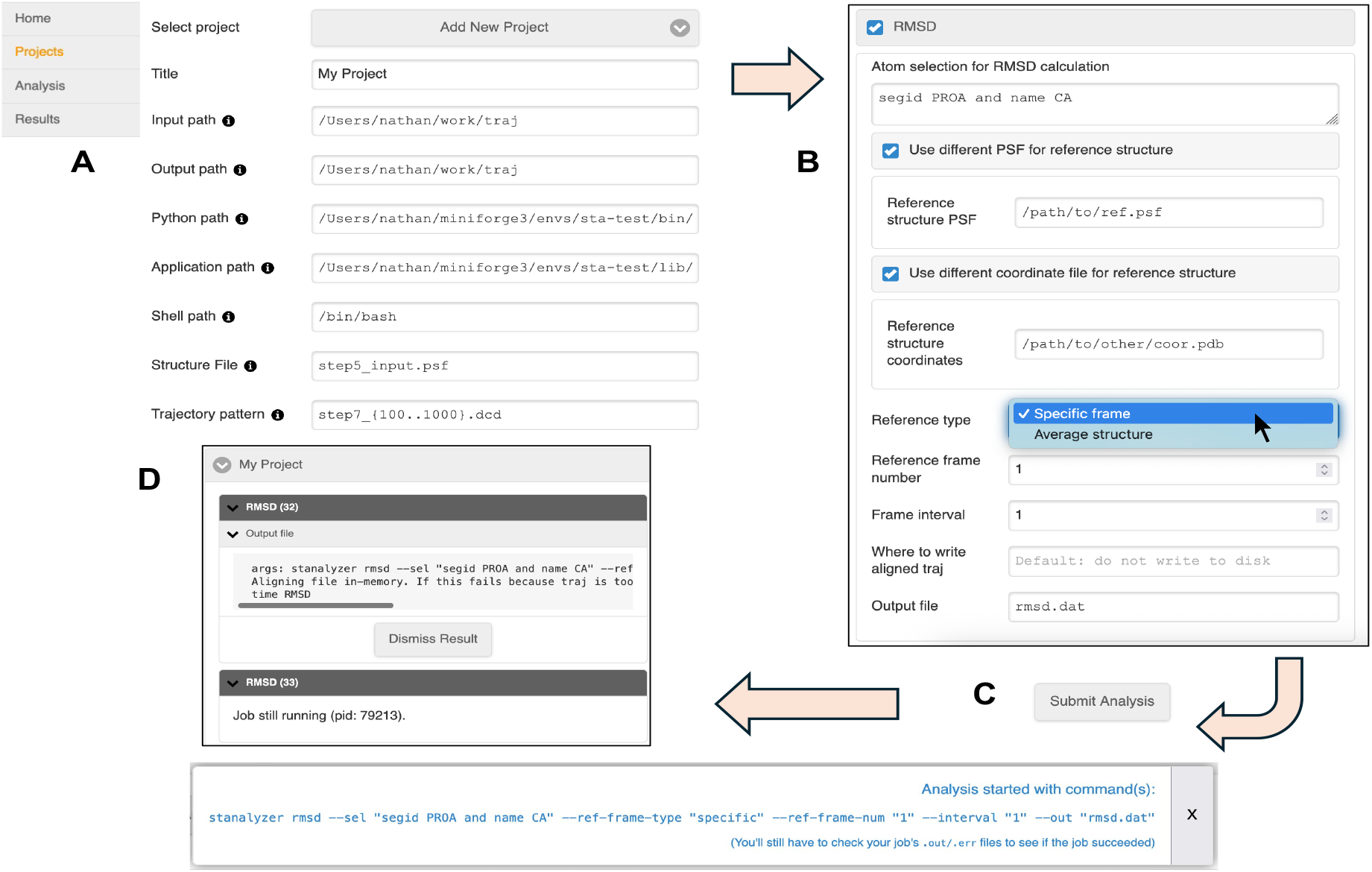
GUI usage example. (A) Users must create a project on the Projects page before any analysis can be run. The main menu is shown on the left. (B) After selecting a project, the analysis page lets users choose a set of programs to run and customize their settings. (C) When users “ Submit” the analysis form, the web server responds with the arguments used for each selected analysis. (D) Each requested analysis has a unique ID, shown in parentheses. Results indicate whether the job is still running. Complete jobs show up to 2000 lines of the output log.

### Implementation

The CLI program and analysis programs are written in pure Python. Analysis programs use MDAnalysis,^5^ NumPy,^9^ SciPy,^10^ and scikit-learn^19^ to query molecular information and calculate results. The web server is written for the FastAPI framework,^17^ which includes the Starlette ASGI (Asynchronous Server Gateway Interface) framework,^20^ Uvicorn ASGI server,^21^ Pydantic data validation library,^22^ and Jinja HTML template engine.^23^ Responsive UI is achieved with jQuery and jQueryMobile. The software is distributed as a conda-forge recipe and includes a README, which can also be found on the official GitHub project page (https://github.com/nk53/stanalyzer).^24^ The README contains instructions for installation, basic usage, and troubleshooting.

### Applications

We next demonstrate the applications of ST-Analyzer to the analysis of the previously studied biological membranes^25^ and SARS-Cov2 spike protein complex systems.^26^

### Biological membranes

Biological membranes exhibit diverse lipid compositions across species (mammals, plants, and fungi) and organelles (plasma membrane, endoplasmic reticulum, and Golgi apparatus).^27,28^ Quantitative analysis of lipid properties such as area per lipid (APL) and deuterium order parameter (*S*_CD_) is often challenging due to their compositional complexity and the heterogeneous lipid distributions between leaflets (**Figure 3A** for example).

**Figure 3.**
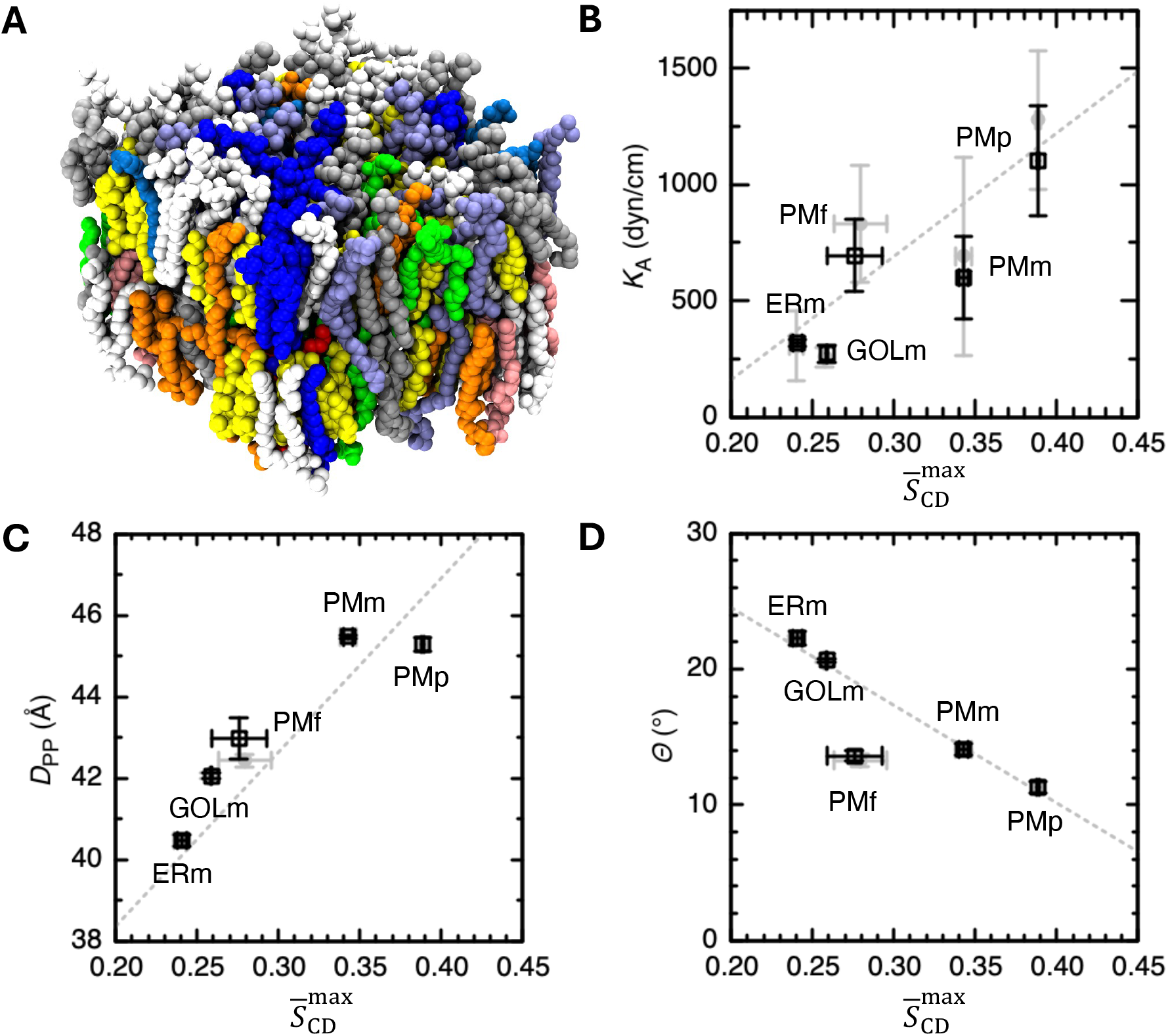
(A) Snapshot of a mammalian plasma membrane. Lipid heavy atoms are shown as spheres, colored by lipid type. (B-D) Correlation of maximum acyl chain order parameter 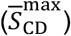 with (B) area compressibility modulus (*K*_A_), (C) membrane thickness (*D*_PP_), and (D) sterol tilt angle (*Θ*). The results obtained using ST-Analyzer are shown as black open squares, while the results from previous analyses^25^ are shown as filled grey circles. Error bars indicate 90% confidence intervals (CI = 2.92 × standard error; *n* = 3). Grey dotted lines show the linear fits from the previous analysis of 16 multicomponent and 3 single-component membrane systems.

Using the membrane models from the previous simulation study,^25^ we analyzed five membrane systems: mammalian plasma (PMm), plant plasma (PMp), fungal plasma (PMf), mammalian ER (ERm), and Golgi (GOLm) membranes. For each system, ST-Analyzer was used to calculate membrane properties including the area compressibility modulus (*K*_A_), membrane thickness (*D*_PP_), sterol tilt angle (*Θ*), APL, and *S*_CD_ (interested readers are referred to SI **S3. Membrane Analyses** for details). For each membrane, the analyses were performed on the final 500 ns of three independent 1-μs simulations, and the results are reported as mean values with 90% confidence intervals.

**Figures 3B-D** illustrate the relationship between *K*_A_, *D*_PP_, *Θ*, and 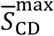. Here,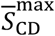 is defined as the maximum *S*_CD_ obtained from the 16:0 *sn*-1 chain, averaged over all phospho- and lyso-lipids in each membrane (**Figure S4**). Our results, including *S*_CD_ profiles (**Figure S4**), APLs (**Tables S1-5**), and membrane thicknesses (**Table S6**) obtained using ST-analyzer are in good agreement with previously reported analyses.

### Spike protein

The other test system is the SARS-CoV-2 spike protein in complex with an antibody targeting the receptor binding domain (RBD).^25^ The trimeric spike protein is anchored in the viral envelope and mediates viral attachment to the host receptor. The RBD adopts open and closed conformations, with the open state being less buried, thereby permitting direct engagement with the host receptor. However, many neutralizing antibodies (NAbs) can bind the RBD in either state, depending on the spatial occupancy of surrounding protein residues and glycans.^29–32^ In this test system, the spike protein contains two protomers in the open state, both of which are bound by the antibody named C105 (**Figure 4A**).^32^

**Figure 4.**
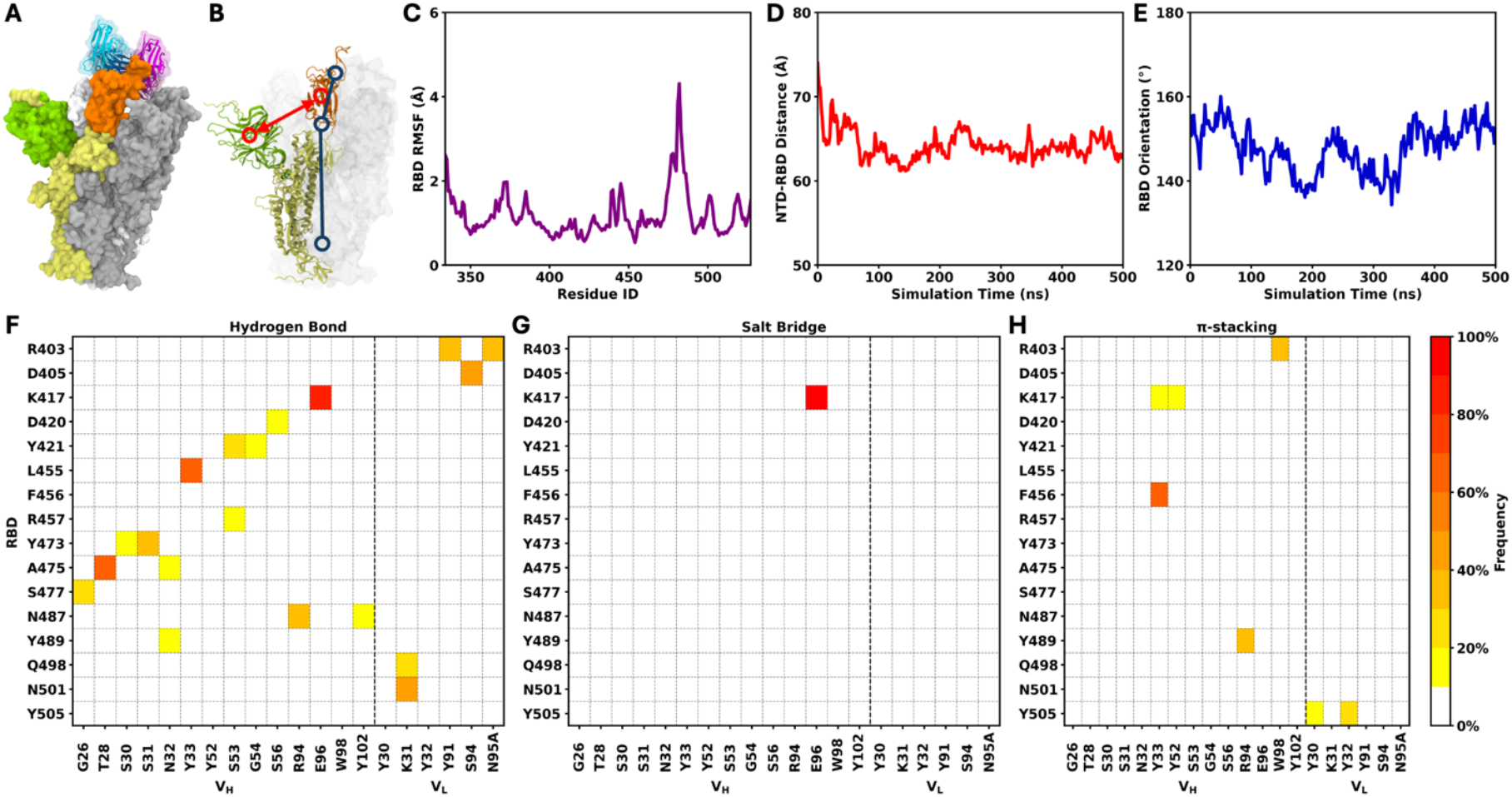
SARS-CoV-2 spike protein. (A) Spike protein in complex with the antibody C105. The three protomers of the spike protein are shown in yellow, gray, and white. The RBD and NTD in the first protomer (yellow) are highlighted in orange and green, respectively. The antibodies bound to two open-state RBDs are shown in blue and magenta. (B) Illustration of the NTD-RBD distance (red) and RBD orientation (blue). (C) RMSF of the RBD. (D) NTD-RBD distance and (E) RBD orientation as functions of simulation time. (F–H) Frequencies of residue pairs forming hydrogen bonds, salt bridges, and Π-stacking interactions between the RBD of the first protomer and the bound antibody (orange and blue in panel A).

We analyzed a 500-ns simulation trajectory consisting of 500 frames from our previous study^26^ and performed the following analyses using ST-Analyzer: system size, secondary structure, RMSD, RMSF, bond statistics, hydrogen bond, salt bridge, π-stacking interaction, water bridge and contact residence time. The system size analysis outputs the box dimension in each frame of the simulation trajectory, which is approximately 236×236×236 Å^3^ throughout the trajectory (**Figure S5**). The secondary structure analysis outputs the DSSP-encoded^33,34^ secondary structure assignment for each residue in each frame (**Figure S6**). The RMSD analysis outputs the structural deviation of the selected protein region over time (**Figure S7**), while the RMSF analysis outputs per-residue fluctuations (**Figure S8**). As an example, the RMSF profile of the RBD in the first protomer of the spike protein is shown in **Figure 4C**, which is consistent with our previous work. The bond statistics analysis quantifies not only bond lengths, angles, and dihedrals formed by individual atoms, but also distances, angles, and dihedrals defined between selected groups of atoms. We calculated the distance between the centers of mass (COMs) of the N-terminal domain (NTD) and the RBD, as well as the angle defined by two atom groups located at both ends of the RBD and one atom group located in the center of the protein stem (**Figure 4B**), and the results are shown in **Figures 4D, E** and **Figure S9**. The outputs of hydrogen bond and water bridge analyses are sorted by frame indices, with each entry consisting of the frame index and the atoms forming the hydrogen bond/water bridge, as well as the bond length and angle in hydrogen bond analysis (**Figures S10-11**). The outputs of salt bridge and π-stacking interaction analyses are sorted by the interaction frequency, with each entry consisting of the interacting residue pair and the indices of frames in which the interaction occurs (**Figures S12-13**). The frequencies of hydrogen bond, salt bridge, and π-stacking interactions between the RBD of the first protomer and the bound antibody are shown in **Figures 4F-H**. When these interaction types are considered together, the resulting interaction pattern is similar to that observed in our previous work. The contact residence time analysis outputs the interacting residue pairs along with the mean and standard deviation of their contacting durations, quantified by the number of frames (**Figure S14**).

Together with the streamlined setup by its Web GUI, these results demonstrate ST-Analyzer as a practical tool for quantitative analysis of complex biological membrane properties and protein-antibody interactions.

### Comparison with Other Software

Popular simulation packages such as GROMACS,^2^ NAMD/VMD,^3,35,36^ and CHARMM^4^ contain extensive simulation, model manipulation, format conversion, and analysis capabilities, but users are still responsible for ensuring the soundness of their simulation/analysis methodology, syncing files between workspace and simulation computers, and ensuring that each step of simulation/analysis is run in the appropriate order. In practice, this is accomplished with ad hoc scripts that must be adapted to each project. Because these software packages are open source, anyone can contribute an analysis plugin to GROMACS, VMD, or CHARMM, but doing so requires understanding implementation details of each program. GROMACS, NAMD, and VMD are written mainly in C/C++; VMD offers a Tcl scripting interface in its pre-built binaries and can be compiled with Python scripting support; CHARMM is implemented in Fortran and comes with its own scripting language. Recently, GROMACS and CHARMM have added Python APIs to access their internal functions.^37–39^ Commercial packages such as Material Studio,^40^ Maestro,^41^ and Amsterdam Modeling Suite^42^ have streamlined workflows, but their proprietary nature impedes the sharing of workflows with the academic community. Other open source packages such as MDAnalysis,^5^ MDTraj,^6^ PyTraj,^7^ and PyLOOS^8^ emphasize the ability for users to create and share analysis routines.

Researchers commonly use many different simulation and analysis packages to handle different aspects of simulation and analysis, depending on what analysis routines that they have on hand and which frameworks that they are familiar with. Rather than supersede any of these packages, ST-Analyzer aims to be a simulation platform-agnostic framework that can encapsulate analysis workflows or be used by them. In this context, a main goal of our implementation is to allow experienced users to incorporate new routines with minimal knowledge of the framework’s implementation details, relative to other frameworks.

Because PC users must open a terminal, activate a conda environment, then issue the *sta-server* command, starting the ST-Analyzer application is not yet as seamless as clicking an application icon. Since conda is a popular package manager and the steps are similar for other conda packages, we anticipate this will not be too much of a barrier for the time being. GUI applications are generally more intuitive than CLIs. While a native desktop application can be designed to use a computer’s resources more efficiently than a locally hosted web GUI, the advantage of a web application is that it can benefit from mature web browsers that value cross-platform usability and backwards compatibility. In effect, updates to the web UI are less subject to the decisions of a single operating system. In principle, only the server host needs any specific combination of software.

### Future Work

At the current stage, ST-Analyzer is a minimum viable product that can be useful to researchers as-is. However, there are clear opportunities for improvement. The milestones below are listed approximately in order of importance.

#### User account management

Because of ST-Analyzer’s responsive UI, if the server is configured to accept remote connections, the user can even view and submit analyses from a mobile browser. Although we verified mobile usability, the current lack of a user account creation and management system renders this operation insecure except on an encrypted single-user network or configuring certificate-based authentication for both server and client machines—a tedious process. Thus, we do not encourage enabling remote access until a future update enables multiple user accounts.

#### Plugins menu

While the procedure for adding a custom analysis to ST-Analyzer’s CLI is straightforward (see **Figure S2**), it still lacks a standard mechanism to package, share, and install it. Instead, each user currently must manually place their code within their conda environment’s site-packages/stanalyzer/analysis subdirectory. There is currently no way to add analysis to the GUI without manually editing internal ST-Analyzer configuration files. To formalize the plugin system, we plan to (1) separate plugin GUI configuration from the built-in GUI, (2) link documentation of the plugin development process in the app’s main menu, and (3) enable plugin installation and removal from the GUI.

#### Results visualization

The current results page only allows checking the status of an analysis task. The most obvious thing to do after generating a data file is to plot it. While generating publication-ready graphs is outside the intended scope of ST-Analyzer, we expect that users would still benefit from the ability to generate standard plots within the GUI.

#### Remote job submission

The original intent of ST-Analyzer included the ability to remotely submit jobs to a scheduler on behalf of the user. We anticipate that the diverse range of security and usability requirements will require careful consideration before its own dedicated update.

## Conclusions

ST-Analyzer consists of four fundamental components: (1) a set of configurable analysis programs, (2) a CLI for the programs that handles reading and writing common default settings for different analysis projects, (3) a GUI that facilitates project settings management, selecting individual analysis settings, invoking the CLI, and checking job status, and (4) a web server that tracks job status and mediates communication between GUI and CLI.

Although there was a previous version of ST-Analyzer with the above capabilities, the difficulty of installation made it clear that a redesign was necessary. This updated version greatly simplifies the installation, update, and software development processes by contributing it as a package to the conda-forge repository. Recipes for conda packages are robust to shifting software ecosystems because they describe the OS and software version requirements necessary to install and run the package. The conda-forge distribution framework’s continuous integration services streamline the testing and update processes, enabling users to receive hotfixes and general updates with minimal effort. ST-Analyzer requires additional future work for the GUI to fully support plugin management, multi-user workstations, and secure networking. However, we still anticipate that the available ST-Analyzer analysis programs and their Python implementations will be useful to experts for quickly setting up common analysis tasks and to nonexperts as a guided introduction to simulation analysis using both GUI and CLI.

## Acknowledgments

This work is supported by NIH R35 GM153458. The authors thank to all Im lab members who contributed to the ST-Analyzer project.

## Data and Software Availability

The ST-Analyzer source code is freely available at https://github.com/nk53/stanalyzer. The README includes instructions for development installation, which allows one to see the effect of source changes without having to manage conda re-packaging. The software is licensed under the MIT License, which allows unrestricted use for academic, commercial, or personal purposes.

