## Supporting Information for "ST-Analyzer: A Packaged Web and Command-Line Interface for Simulation Trajectory Analysis"

### S1. Schema for `project.json`

The example in **Figure S1** shows all required keys, though the files created by ST-Analyzer may contain other optional fields. In the description below, an asterisk (*) marks items are auto-filled with default values obtained from ST-Analyzer's runtime environment when on the GUI's Project page or the CLI's config utility. Items marked with a cross (†) are currently unused and will either be used in a future update or removed. A double cross (‡) implies both * and †.

**`title`:** Display name for project in GUI.

**`input_path`:** Absolute path to directory containing PSF and trajectory files.

**`output_path`:** Absolute path to directory where analysis output should be written (maybe the same as input_path).

**`shell_path`:** * Absolute path to shell that the GUI should use to spawn the CLI.

**`application_path`:** ‡ Absolute path to GUI application.

**`python_path`:** ‡ Absolute path to the version of Python running ST-Analyzer.

**`psf`:** Path to PSF file, relative to `input_path`.

**`traj`:** Path to trajectory files, relative to `input_path`. Shell globbing and brace expansion may be used to select multiple files.

**`scheduler`:** † Currently, the only valid option is interactive, but a future update may enable job delegation via SLURM or PBS.

**`time_step`:** A number followed by one of [ms, µs, us, ns, ps, fs]. Used by some analyses to increment the time column of an output.

### S2. Schema for `analysis.yml`

The example in Figure S3 shows the configuration that yields the web form shown in Figure 2B. The outermost key must match the stem of the analysis program being described; thus the key for rmsd.py is rmsd. It must contain the following items:

#### S2.1 `FormSpec`

**`label`:** (string) Display name of analysis in GUI.

**`options`:** (mapping of string to `FieldSpec`) One or more fields for specifying CLI flags.

#### S2.2 `FieldSpec`

**`label`:** (string) Text description of field to put in a `<label>` tag.

**`type`:** (Jinja macro) See `TypeSpec` (**S3.3**).

**`options`:** (mapping of string to string OR `FieldSpec`) If `type` is `select`, then each contained key is the HTML name of the option to generate and its values are that option's display text. Otherwise, `type` is automatically set to `form.checkbox` and options must be a nested subform (`FieldSpec`) whose visibility is toggled by the checkbox.

**`classes`:** (optional list of strings) CSS classes to add to the field for display. Currently, the only class used is `code`, which uses a fixed-width font to display the field's text value.

**`visibility`:** (optional dict of visibility rules) See **S3.4 Visibility Rules**.

**`html_attr ...`:** (Optional) Additional keys are interpreted as HTML attributes that should be set verbatim. Commonly used attributes are `value` (field's default value, if any) and `placeholder` (text to display if/when `value` is empty).

#### S2.3 `TypeSpec`

**`form.checkbox`:** HTML `<input type=checkbox>` element. Set checked to true to make the box checked by default.

**`form.input`:** HTML `<input type=text>` element.

**`form.path`:** A shorthand for `<input type=text class="code">`.

**`form.input_integer`:** HTML `<input type=number>`. On supporting devices, it adds up/down buttons that increase/decrease the integer value by 1. A future update may add client-side input validation so that "submit" will warn if a non-integer value is entered.

**`form.select`:** HTML `<select>` element, with contained `<option>` elements provided by an options dict containing option name: value entries. See `ref_frame_type` in **Figure S3**.

**`form.textarea`:** HTML `<textarea>` element. Basically a resizable text `<input>`.

#### S2.4. `Visibility` Rules

Each entry in `visibility` consists of a rule whose name determines the rule's type and target. **Figure S3** shows one visibility rule: **rmsd_ref_frame_type:** `specific`. The general format for a visibility rule is:

{*rule type*}_{*analysis name*}_{*field name*}: *value*

Two rule types are allowed:

**`allowed`:** The field will only be visible if its value is *value*.

**`disallowed`:** The field will only be visible if its value is NOT *value*.

*analysis name* indicates which analysis form contains the field controlling the current field's visibility. *field name* is the name of the controlling field. To allow multiple analysis programs to provide CLI options with the same name without causing a name conflict on the Analysis page, ST-Analyzer always prepends the name of the analysis to the field name before rendering its HTML. Thus, both *analysis name* and *field name* are required to name another field. In most cases, we expect *analysis name* to be the same as the form being defined, but this allows field visibility to be affected by any input on the same page. Lastly, a field with multiple visibility rules will only be visible if all of the given rules are satisfied.

Thus, the rule **allowed_rmsd_ref_frame_type:** `specific` in `ref_frame_num`'s definition causes `ref_frame_num` to be hidden unless the value of `ref_frame_type` is `specific`.

### S3. Membrane Analyses

This section describes the membrane properties analyzed in this work and the methodologies used to compute the area per lipid (APL) and the deuterium order parameter ($S_{CD}$). All membrane analysis modules in ST-Analyzer assume a planar membrane geometry, which is typical for membrane simulations.

The area compressibility modulus $K_A$ is defined as $K_{\mathrm{A}} = k_{\mathrm{B}}T\langle A\rangle/\langle \delta A^2\rangle$, where $k_B$ is the Boltzmann constant, $T$ is temperature, and $\langle A\rangle$ and $\langle \delta A^2\rangle$ are the average and fluctuation of the simulation box XY-area. The sterol tilt angle $\Theta$ is defined as the angle between the sterol vector (from C17 to C3) and the membrane normal (Z-axis). The membrane thickness $D_{PP}$ was defined as the distance between the average Z positions of phosphorous atoms in opposing leaflets. The hydrophobic thickness $D_{CC}$ was defined analogously using the first aliphatic carbons (C22, C32, C4S, or C2F) or the sterol hydroxyl carbons (C3). Leaflet thicknesses were measured relative to the bilayer midplane ($Z$ = 0). In ST-Analyzer, the midplane is defined by default as the mean Z-positions of representative atoms in the bilayer; optionally, it can be defined as the crossing point of leaflet Z-density profiles. Although local thicknesses can be defined analogously, they are not currently supported.

The APLs for individual lipid types were calculated using a Voronoi tessellation approach.[1] Lipids were represented by selected atoms projected onto the *XY*-plane, and the molecular area was calculated as the sum of corresponding Voronoi cell areas. The APLs for individual types were obtained by averaging over molecules of the same type within each leaflet and, for symmetric bilayers, over both leaflets. Phospholipids were represented by three atoms: the glycerol backbone (C2) and two acyl (C21 and C31) carbons. Sphingolipids were represented by analogous atoms: C2S, C3S, and C1F. Lysolipids and sterols were represented by a single atom, the acyl carbon (C31) and the hydroxyl oxygen (O3), respectively.

The deuterium order parameter was calculated as $S_{\mathrm{CD}} = |\langle 3\cos^2\theta - 1\rangle/2|$, where $\theta$ is the angle between C-H vector and the membrane normal ($Z$-axis). Analysis of $S_{CD}$ in multicomponent membranes is often challenging due to variations in tail number and length across lipid classes (e.g., phospholipids, sphingolipids, and lysolipids, and lipopolysaccharides). ST-Analyzer addresses this challenge by allowing users to specify the first aliphatic carbon in each tail for each lipid type. Remaining tail carbons are automatically identified at the beginning of the analysis. The calculated $S_{CD}$'s were averaged by lipid type and leaflet, and for symmetric bilayers, over both leaflets. In this work, $S_{CD}$ profiles for all lipid types were computed using ST-Analyzer for each membrane. For comparative analysis, the $S_{CD}$ profiles corresponding to the 16:0 *sn*-1 chain were subsequently averaged in a post analysis to evaluate correlations with their maximum value ($\overline{S}_{\mathrm{CD}}^{\mathrm{max}}$) and other membrane properties shown in **Figure 3B-D**.

**Figure S1.** Example `project.json` configuration.

```
    "title": "My Project",
    "input_path": "/Users/nathan/work/traj",
    "output_path": "/Users/nathan/work/traj/out",
    "shell_path": "/bin/bash",
    "application_path": "/Users/nathan/src/stanalyzer/src/stanalyzer",
    "python_path": "/Users/nathan/miniforge3/envs/sta/bin/python",
    "psf": "step5_input.psf",
    "traj": "step7_{1..999}.dcd",
    "scheduler": "interactive",
    "time_step": "0.1 ns"
```

**Figure S2.** Wrapper script for external program.

```
import os
import sys
from stanalyzer.cli import FakeParser

ANALYSIS_NAME = 'wrapper'


def get_parser():
    # FakeParser.parse_args() always returns an empty namespace
    return FakeParser()


def main(settings=None):
    program = 'my_program'

    # replace 'stanalyzer' and 'wrapper' with program name
    args = [program] + sys.argv[2:]

    # if called like `stanalyzer wrapper arg1 arg2`, then the result is:
    #   my_program arg1 arg2
    os.execvp(program, args)


if __name__ == '__main__':
    main()
```

**Figure S3.** Analysis GUI configuration showing most features.

```
rmsd:
  label: RMSD
  options:
    sel:
      label: Atom selection for RMSD calculation
      type: form.textarea
      classes: [code]
      value: segid PROA and name CA
    ref_psf:
      label: Use different PSF for reference structure
      options:
        path:
          label: Reference structure PSF
          type: form.path
    ref_coor:
      label: Use different coordinate file for reference structure
      options:
        path:
          label: Reference structure coordinates
          type: form.path
    ref_frame_type:
      label: Reference type
      type: form.select
      options:
        specific: Specific frame
        average: Average structure
    ref_frame_num:
      label: Reference frame number
      type: form.positive_integer
      value: 1
      visibility:
        allowed_rmsd_ref_frame_type: specific
    interval:
      label: Frame interval
      type: form.positive_integer
    align_out:
      label: Where to write aligned traj
      type: form.path
      value: ""
    out:
      label: Output file
      type: form.path
      value: rmsd.dat
```

**Figure S4.** Deuterium order parameter ($S_{CD}$) of the 16:0 *sn*-1 acyl chain in five membranes (open squares). Shown together as grey filled circles are $S_{CD}$ from the previous analyses.[2] Error bars indicate 90% confidence intervals (CI = 2.92 × standard error; *n* = 3).

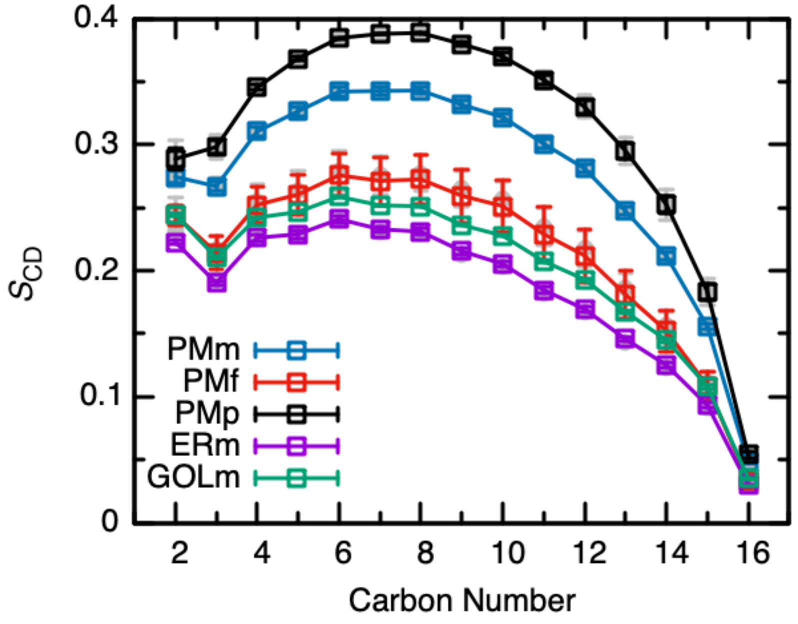

**Figure S5.** The output of system size.

```
#time xtla xtlb xtlc alpha beta gamma volume
1   235.81 235.81 235.81 90.000 90.000 90.000 1.3112e+07
2   235.60 235.60 235.60 90.000 90.000 90.000 1.3077e+07
3   235.63 235.63 235.63 90.000 90.000 90.000 1.3083e+07
... ...
498 235.67 235.67 235.67 90.000 90.000 90.000 1.3089e+07
499 235.57 235.57 235.57 90.000 90.000 90.000 1.3073e+07
500 235.58 235.58 235.58 90.000 90.000 90.000 1.3074e+07
```

**Figure S6.** The output of secondary structure.

```
#DSSP Code  Description
#    H      Alpha-helix
#    B      Beta-bridge (residue in isolated beta-bridge)
#    E      Strand (extended strand, participates in beta-ladder)
#    G      Helix_3 (3_10-helix)
#    I      Helix_5 (pi-helix)
#    P      Helix_PPII (Kappa-helix (poly-proline II helix))
#    T      Turn (hydrogen-bonded turn)
#    S      Bend
#    -      Loop
#residue secondary_structures_in_each_frame
PROA_PHE_318 ------------------------------------------------------------...
PROA_ARG_319 --------------------------------------------P-----P---------...
PROA_VAL_320 P---P-P-PPPP-PPP--PPPP-PP-PPPPP--PPP-PPPPP-PP-PP-PPPPPPP--P-P..
... ...
PROA_VAL_539 -EEE-EE-E-E-EEEEEE--EE--EE--E-EEEEEEEEE--EE-E-EEE--EE-E---EE-..
PROA_ASN_540
BEEEBEEBEBEBEEEEEEBBEEBBEEBBEBEEEEEEEEEEBBEEBEBEEEBBEEBEBBBEEB...
PROA_PHE_541 ------------------------------------------------------------...
```

**Figure S7.** The output of RMSD.

```
#time RMSD
1   0.0000
2   2.78794
3   2.27254
... ...
498 2.93979
499 3.09074
500 2.85328
```

**Figure S8.** The output of RMSF.

```
#residue_indices RMSF
318 3.30169
319 2.99466
320 2.84644
... ...
539 2.01532
540 1.91618
541 1.94919
```

**Figure S9.** The output of bond statistics.

```
#Bond Length (Angstrom)[A1_A2]
75.2420
72.6398
72.3287
... ...
#Bond Angle (Degrees)[A3_A4_X5]
155.5499
152.1876
152.8335
... ...
```

**Figure S10.** The output of hydrogen bond.

```
#frame donor hydrogen acceptor distance angle
1   PROA_THR_323_N PROA_THR_323_HN PROA_CYS_538_O 2.823188 168.085044
1   PROA_ILE_326_N PROA_ILE_326_HN PROA_ASN_540_O 2.921437 153.473301
1   PROA_VAL_327_N PROA_VAL_327_HN PROA_THR_531_OG1 2.936248 162.591331
... ...
2   PROA_ILE_326_N PROA_ILE_326_HN PROA_ASN_540_O 2.747978 153.925236
2   PROA_VAL_327_N PROA_VAL_327_HN PROA_THR_531_OG1 2.801376 168.767483
2   PROA_ARG_328_NE PROA_ARG_328_HE PROA_VAL_327_O 2.638863 166.496263
... ...
500 PROE_THR_110_OG1 PROE_THR_110_HG1 PROE_GLU_86_OE1 2.898623 169.919788
500 PROE_VAL_111_N PROE_VAL_111_HN PROE_GLU_86_OE1 3.160522 155.724966
500 PROE_LEU_117_N PROE_LEU_117_HN PROE_GLY_13_O 2.764240 169.753699
```

**Figure S11.** The output of water bridge.

```
#frame site1 water_to_site1 water_to_site2 site2 (each site and water are in
the form of [acceptor, None] or [donor_hydrogen, donor_heavy])
1   [PROA_LYS_417_HN PROA_LYS_417_N] [SOLV_TIP3_214694_OH2 None]
[SOLV_TIP3_214694_H2 SOLV_TIP3_214694_OH2] [PROD_TYR_58_OH None]
1.  [PROA_TYR_421_HH PROA_TYR_421_OH] [SOLV_TIP3_211475_OH2 None]
[SOLV_TIP3_211475_H1 SOLV_TIP3_211475_OH2] [PROD_SER_56_OG None]
1   [PROA_TYR_421_HH PROA_TYR_421_OH] [SOLV_TIP3_211475_OH2 None]
[SOLV_TIP3_211475_OH2 None] [PROD_SER_56_HN PROD_SER_56_N]
... ...
2.  [PROA_TYR_453_HH PROA_TYR_453_OH] [SOLV_TIP3_27970_OH2 None]
[SOLV_TIP3_27970_H1 SOLV_TIP3_27970_OH2] [PROD_GLU_102_OE1 None]
2.  [PROA_SER_477_HN PROA_SER_477_N] [SOLV_TIP3_191760_OH2 None]
[SOLV_TIP3_191760_H1 SOLV_TIP3_191760_OH2] [PROD_GLY_26_O None]
2   [PROA_TYR_505_HH PROA_TYR_505_OH] [SOLV_TIP3_192169_OH2 None]
[SOLV_TIP3_192169_H1 SOLV_TIP3_192169_OH2] [PROE_TYR_35_OH None]
... ...
500 [PROA_GLU_406_OE2 None] [SOLV_TIP3_332237_H2 SOLV_TIP3_332237_OH2]
[SOLV_TIP3_332237_OH2 None] [PROD_TRP_101_HE1 PROD_TRP_101_NE1]
500 [PROA_SER_494_O None] [SOLV_TIP3_350482_H1 SOLV_TIP3_350482_OH2]
[SOLV_TIP3_350482_H2 SOLV_TIP3_350482_OH2] [PROD_GLU_102_OE1 None]
500 [PROA_ASP_405_OD1 None] [SOLV_TIP3_373972_H1 SOLV_TIP3_373972_OH2]
[SOLV_TIP3_373972_H2 SOLV_TIP3_373972_OH2] [PROE_GLY_96_O None]
```

**Figure S12.** The output of salt bridge.

```
#residue1 residue2 frames
PROA_ASP_467 PROA_ARG_457 1 2 3 4 5 6 7 8 9 10 11 12 13 14 15 16 17 18 19...
PROA_ASP_85  PROA_ARG_64  2 3 4 5 6 7 8 9 10 11 12 13 14 15 16 17 18 19
20...
PROA_GLU_406 PROA_ARG_403 1 2 3 4 5 6 7 8 9 10 11 12 13 14 15 16 17 18 19...
... ...
PROA_GLU_406 PROA_ARG_408 291
PROA_GLU_324 PROA_ARG_328 413
PROA_ASP_398 PROA_ARG_466 443
```

**Figure S13.** The output of π-stacking interaction.

```
#residue1 residue2 frames
PROA_TRP_353 PROA_TYR_423 1 2 3 4 5 6 7 8 9 10 11 12 13 14 15 16 17 18 19...
PROA_PHE_392 PROA_PHE_515 1 2 3 4 5 6 7 8 9 10 11 12 13 14 15 16 17 18 19...
PROA_PHE_400 PROA_TYR_423 1 2 3 4 5 6 7 8 9 10 11 12 13 14 15 16 17 18 19...
... ...
PROA_PHE_377 PROA_TYR_365 375
PROA_TYR_52  PROE_LYS_56  392
PROA_TRP_101 PROA_LYS_417 495
```

**Figure S14.** The output of contact residence time.

```
#residue1_resname residue1_ID residue2_resname residue2_ID mean_time std
GLN 3   LEU 4   1.0000 0.0000
ARG 319 GLN 321 1.4348 0.7704
ASN 334 LEU 335 4.4675 6.1993
SER 438 ASN 439 15.828 29.191
ASP 420 TYR 421 70.571 73.793
GLN 506 PRO 507 249.50 36.500
ILE 358 VAL 395 500.00 0.0000
... ...
```

**Table S1.** Lipid compositions and APL of individual lipid types in asymmetric mammalian plasma membrane (PMm).

| Lipid Name | Lipid Head/Tail | # Lipids | | APL (Å$^2$)[a] | | Charge ($e$) |
|---|---|---|---|---|---|---|
| | | Outer | Inner | Outer | Inner | |
| POPC | PC(16:0/18:1(9Z)) | 16 | 7 | 56.9 (0.7) / 56.9 | 57.1 (0.4) / 57.1 | 0 |
| PLPC | PC(16:0/18:2(9Z,12Z)) | 22 | 11 | 57.3 (0.3) / 57.3 | 57.3 (0.4) / 57.4 | 0 |
| PAPE | PE(16:0/24:4(5Z,8Z,11Z,14Z)) | 3 | 12 | 57.6 (0.7) / 57.6 | 58.4 (0.3) / 58.4 | 0 |
| POPE | PE(16:0/18:1(9Z)) | 3 | 14 | 55.1 (2.7) / 55.1 | 56.6 (0.6) / 56.6 | 0 |
| POPI | PI(16:0/18:1(9Z)) | 0 | 5 | – / – | 56.8 (1.0) / 56.8 | -1 |
| PAPS | PS(16:0/20:4(5Z,8Z,11Z,14Z)) | 0 | 11 | – / – | 58.3 (0.6) / 58.3 | -1 |
| POPA | PA(16:0/18:1(9Z)) | 0 | 1 | – / – | 55.7 (2.3) / 55.7 | -1 |
| SSM | SM(d18:1/18:0) | 11 | 5 | 50.1 (1.2) / 50.1 | 49.7 (1.0) / 49.7 | 0 |
| NSM | SM(d18:1/24:1) | 11 | 5 | 49.5 (0.6) / 49.5 | 49.9 (1.4) / 49.9 | 0 |
| CMH | GlcCer(d18:1/16:0) | 4 | 0 | 47.3(0.7) / 47.3 | – / – | 0 |
| CHOL | Cholesterol | 37 | 29 | 27.9 (0.2) / 27.9 | 28.7 (0.5) / 28.7 | 0 |
| Total | | 107 | 100 | | | |

[a]Leaflets were assigned at every frame in this work. 90% confidence interval (CI = 2.92 × standard error; $n$ = 3) are shown in parentheses. The results from the previous simulation study[2] are preceded by a slash.

**Table S2.** Lipid compositions and APL of individual lipid types in asymmetric fungal plasma membrane (PMf).

| Lipid Name | Lipid Head/Tail | # Lipids | | APL (Å$^2$)[a] | | Charge ($e$) |
|---|---|---|---|---|---|---|
| | | Outer | Inner | Outer | Inner | |
| DYPC | PC(16:1(9Z)/16:1(9Z)) | 8 | 10 | 57.0 (1.7) / 56.1 | 62.8 (2.8) / 62.2 | 0 |
| YOPC | PC(16:1(9Z)/18:1(9Z)) | 3 | 6 | 53.7 (2.6) / 51.9 | 61.7 (1.3) / 61.7 | 0 |
| POPE | PE(16:0/18:1(9Z)) | 3 | 10 | 52.9 (1.4) / 53.4 | 61.0 (2.0) / 60.8 | 0 |
| PYPE | PE(16:0/16:1(9Z)) | 1 | 5 | 51.4 (5.6) / 51.9 | 60.1 (2.7) / 59.8 | 0 |
| YOPE | PE(16:1(9Z)/18:1(9Z)) | 3 | 6 | 53.9 (4.6) / 54.7 | 60.9 (2.2) / 60.7 | 0 |
| POPI | PI(16:0/18:1(9Z)) | 5 | 20 | 55.1 (1.5) / 55.3 | 61.1 (2.1) / 60.4 | -1 |
| POPS | PS(16:0/18:1(9Z)) | 7 | 31 | 56.7 (1.4) / 57.5 | 59.6 (1.9) / 59.6 | -1 |
| YOPA | PA(16:1(9Z)/18:1(9Z)) | 3 | 8 | 52.7 (8.2) / 54.0 | 60.3 (2.1) / 59.9 | -1 |
| ERG | Ergosterol | 49 | 4 | 26.6 (0.8) / 26.5 | 30.1 (1.4) / 29.6 | 0 |
| MIPC | MIPC(d18:1/16:0) | 54 | 0 | 50.8 (0.6) / 50.3 | – / – | 0 |
| Total | | 136 | 100 | | | |

[a]Leaflets were assigned at every frame in this work. 90% confidence interval (CI = 2.92 × standard error; $n$ = 3) are shown in parentheses. The results from the previous simulation study[2] are preceded by a slash.

**Table S3.** Lipid compositions and APL of individual lipid types in asymmetric plant plasma membrane (PMp).

| Lipid Name | Lipid Head/Tail | # Lipids | | APL (Å$^2$)[a] | | Charge ($e$) |
|---|---|---|---|---|---|---|
| | | Outer | Inner | Outer | Inner | |
| DPPC | PC(16:0/16:0) | 3 | 6 | 57.5 (0.5) / 57.6 | 54.3 (0.5) / 54.3 | 0 |
| LLPC | PC(18:2(9Z,12Z)/18:3(9Z,12Z,15Z)) | 7 | 10 | 59.4 (0.3) / 59.4 | 58.3 (0.4) / 59.3 | 0 |
| SOPC | PC(18:0/18:1(9Z)) | 3 | 4 | 58.7 (0.7) / 58.7 | 57.1 (1.2) / 57.1 | 0 |
| DPPE | PC(16:0/16:0) | 8 | 12 | 55.0 (0.4) / 55.0 | 53.9 (0.3) / 53.9 | 0 |
| LLPE | PE(18:2(9Z,12Z)/18:3(9Z,12Z,15Z)) | 7 | 10 | 60.1 (0.5) / 60.1 | 57.3 (0.3) / 57.3 | 0 |
| SOPE | PE(18:0/18:1(9Z)) | 1 | 2 | 56.5 (1.4) / 56.5 | 55.7 (0.5) / 55.7 | 0 |
| DPPA | PA(16:0/16:0) | 2 | 4 | 54.2 (0.6) / 54.2 | 53.7 (0.2) / 53.7 | -1 |
| LLPA | PA(18:2(9Z,12Z)/18:3(9Z,12Z,15Z)) | 3 | 5 | 60.5 (1.7) / 60.5 | 58.1 (0.1) / 58.1 | -1 |
| SOPA | PA(18:0/18:1(9Z)) | 0 | 2 | – / – | 56.5 (0.9) / 56.5 | -1 |
| DPPI | PI(16:0/16:0) | 0 | 3 | – / – | 55.2 (0.3) / 55.2 | -1 |
| LLPI | PI(18:2(9Z,12Z)/18:3(9Z,12Z,15Z)) | 0 | 2 | – / – | 57.7 (1.8) / 57.7 | -1 |
| LLPS | PS(18:2(9Z,12Z)/18:3(9Z,12Z,15Z)) | 1 | 2 | 59.3 (1.9) / 59.3 | 58.1 (0.9) / 58.1 | -1 |
| DPPG | PG(16:0/16:0) | 0 | 4 | – / – | 56.1 (0.4) / 56.1 | -1 |
| DGDG | DGDG(18:3(9Z,12Z,15Z)/18L3(9Z,12Z,15Z)) | 0 | 2 | – / – | 59.4 (1.3) / 59.4 | 0 |
| CMH | GlcCer(d18:1/16:0) | 15 | 0 | 48.3 (0.6) / 48.3 | – / – | 0 |
| SITO | β-Sitosterol | 33 | 16 | 30.8 (0.1) / 30.8 | 28.9 (0.2) / 28.9 | 0 |
| STIG | Stigmasterol | 23 | 12 | 31.0 (0.3) / 31.0 | 28.4 (0.2) / 28.4 | 0 |
| CAMP | Campesterol | 8 | 4 | 32.2 (0.9) / 32.2 | 28.2 (0.4) / 28.2 | 0 |
| Total | | 114 | 100 | | | |

[a]Leaflets were assigned at every frame in this work. 90% confidence interval (CI = 2.92 × standard error; $n$ =3) are shown in parentheses. The results from the previous simulation study[2] are preceded by a slash.

**Table S4.** Lipid compositions and APL of individual lipid types in mammalian endoplasmic reticulum membrane (ERm).

| Lipid Name | Lipid Head/Tail | # Lipids | APL (Å$^2$)[a] | Charge ($e$) |
|---|---|---|---|---|
| | | Outer or Inner | Outer or Inner | |
| POPC | PC(16:0/18:1(9Z)) | 16 | 63.4 (0.2) / 63.2 | 0 |
| PLPC | PC(16:0/18:2(9Z,12Z)) | 18 | 63.7 (0.4) / 63.5 | 0 |
| SAPC | PE(18:0/20:4(5Z,8Z,11Z,14Z)) | 27 | 65.0 (0.3) / 64.8 | 0 |
| POPE | PE(16:0/18:1(9Z)) | 5 | 62.6 (0.3) / 62.4 | 0 |
| PSPE | PE(16:0/18:0) | 8 | 61.5 (0.8) / 61.3 | 0 |
| SAPE | PE(18:0/20:4(5Z,8Z,11Z,14Z)) | 7 | 64.2 (0.1) / 64.0 | 0 |
| SAPI | PI(18:0/20:4(5Z,8Z,11Z,14Z)) | 3 | 64.3 (0.9) / 64.1 | -1 |
| SLPI | PI(18:0/18:2(9Z,12Z)) | 3 | 62.8 (0.9) / 62.5 | -1 |
| OLPS | PS(18:1(9Z)/18:2(9Z,12Z)) | 3 | 62.6 (0.5) / 60.7 | -1 |
| PSM | SM(d18:1/16:0) | 4 | 54.7 (0.9) / 59.1 | 0 |
| POPA | PA(16:0/18:1(9Z)) | 1 | 62.3 (1.7) / 62.1 | -1 |
| CHOL | Cholesterol | 5 | 29.9 (0.7) / 29.8 | 0 |
| Total | | 100 | | |

[a]Leaflets were assigned at every frame in this work. 90% confidence interval (CI = 2.92 × standard error; $n$ =3) are shown in parentheses. The results from the previous simulation study[2] are preceded by a slash.

**Table S5.** Lipid compositions and APL of individual lipid types in mammalian Golgi membrane (GOLm).

| Lipid Name | Lipid Head/Tail | # Lipids | APL (Å$^2$)[a] | Charge (*e*) |
|---|---|---|---|---|
| | | Outer or Inner | Outer or Inner | |
| POPC | PC(16:0/18:1(9Z)) | 11 | 61.9 (0.3) / 61.0 | 0 |
| PLPC | PC(16:0/18:2(9Z,12Z)) | 14 | 62.1 (0.3) / 61.2 | 0 |
| SAPC | PC(18:0/20:4(5Z,8Z,11Z,14Z)) | 20 | 63.5 (0.4) / 62.6 | 0 |
| PLPE | PE(16:0/18:2(9Z,12Z)) | 4 | 60.8 (1.4) / 60.1 | 0 |
| PSPE | PE(16:0/18:0) | 8 | 59.2 (1.0) / 58.4 | 0 |
| SAPE | PE(18:0/20:4(5Z,8Z,11Z,14Z)) | 5 | 62.6 (1.0) / 61.8 | 0 |
| PSPI | PI(18:0/18:0) | 2 | 59.8 (0.6) / 58.9 | -1 |
| POPI | PI(18:0/18:1(9Z)) | 5 | 60.3 (0.7) / 59.3 | -1 |
| SAPI | PS(18:0/20:4(5Z,8Z,11Z,14Z)) | 2 | 63.9 (2.6) / 62.9 | -1 |
| POPS | PS(16:0/18:1(9Z)) | 4 | 60.5 (0.8) / 59.6 | -1 |
| TSM | SM(d18:0/22:0) | 12 | 54.5 (0.3) / 57.4 | 0 |
| LPC16[b] | PC(16:0/0:0) | 5 | 34.2 (0.3) / 41.4 | 0 |
| CHOL | Cholesterol | 8 | 28.8 (0.6) / 28.2 | 0 |
| Total | | 100 | | |

[a]Leaflets were assigned at every frame in this work. 90% confidence interval (2.92 × standard error; *n* = 3) are shown in parentheses. The results from the previous simulation study are preceded by a slash. [b]In this work, LPC16 was represented using a single atom (C31) for Voronoi tessellation, while two atoms (C2 and C31) were used in the previous analyses.[2]

**Table S6.** Bilayer ($D_{PP}$) and hydrophobic ($D_{CC}$) thicknesses for five membranes.

| Membrane | Thickness (Å)[a] | |
|---|---|---|
| | $D_{PP}$[b] | $D_{CC}$[c] |
| PMm | 45.5 (0.0) / 45.5 (0.1) | 34.4 (0.0) / 34.5 (0.0) |
| PMf | 43.0 (0.5) / 42.2 (0.1) | 32.0 (0.5) / 31.9 (0.1) |
| PMp | 45.3 (0.2) / 45.3 (0.1) | 34.2 (0.1) / 34.2 (0.0) |
| ERm | 40.5 (0.1) / 40.2 (0.1) | 29.5 (0.1) / 29.3 (0.1) |
| GOLm | 42.1 (0.1) / 41.9 (0.1) | 31.0 (0.1) / 30.9 (0.1) |

[a]90% confidence interval (2.92 × standard error; $n$ = 3) are shown in parentheses. The results from previous analyses,[2] with standard errors in parentheses are preceded by a slash. [b]$D_{PP}$ (bilayer thickness) as the distance between the average Z positions of phosphorous atoms (or phosphate groups) in opposing leaflets. [c]$D_{CC}$ (hydrophobic thickness) as the distance between average Z positions for the first aliphatic carbons in lipid tails (C22, C32, C4S, or C2F) or the hydroxyl carbon (C3) in sterols in opposing leaflets.
